# Evaluating CRISPR-based Prime Editing for cancer modeling and CFTR repair in intestinal organoids

**DOI:** 10.1101/2020.10.05.325837

**Authors:** Maarten H. Geurts, Eyleen de Poel, Cayetano Pleguezuelos-Manzano, Léo Carrillo, Amanda Andersson-Rolf, Matteo Boretto, Jeffrey M. Beekman, Hans Clevers

**Affiliations:** Hubrecht Institute, Royal Netherlands Academy of Arts and Sciences (KNAW) and University Medical Center Utrecht, 3584 CT Utrecht, the Netherlands; Oncode Institute, Hubrecht Institute, 3584 CT Utrecht, the Netherlands; Department of Pediatric Respiratory Medicine, Wilhelmina Children’s Hospital, University Medical Center, Utrecht University, 3584 EA Utrecht, the Netherlands; Regenerative Medicine Utrecht, University Medical Center, Utrecht University, 3584 CT Utrecht, the Netherlands

## Abstract

Prime editing is a recently reported genome editing tool employing a nickase-cas9 fused to a reverse transcriptase that directly synthesizes the desired edit at the target site. The technique holds great promise for clinical application due to its versatility. Here, we explore the use of prime editing in human intestinal organoids. Common TP53 mutations were modeled in human adult stem cell with notable efficiency differences. Next, we functionally repaired the cystic fibrosis CFTR-F508del mutation and compared prime editing to CRISPR/Cas9-mediated homology directed repair and adenine base editing on the CFTR-R785* mutation. Despite encountering varying editing efficiencies and undesired mutations, these results underline the broad applicability of prime editing for modeling oncogenic mutations and showcase the potential clinical application of this technique, pending further optimization.

## Main text

The field of genome engineering has been revolutionized by the development of the efficient genome editing tool CRISPR/Cas9. In CRISPR/Cas9-mediated genome engineering, the effector protein cas9 is guided towards the target site in the genome by an RNA guide (Jinek et al., 2012). Upon target recognition, cas9 generates a double stranded break (DSB) that can be exploited for a variety of genome engineering strategies (Cong et al., 2013; Mali et al., 2013). Due to the easy reprogrammability and high efficiency of CRISPR/Cas9, the technology is widely used for gene modification and is considered to be the most promising tool for clinical gene editing. However, the repair of DSBs is often error-prone and can result in unwanted DNA damage at the target site as well as at off-target sites that closely resemble the guide RNA (Cho et al., 2014; Fu et al., 2013; Kosicki et al., 2018; Pattanayak et al., 2013). These issues have been circumvented by the development of Cas9 fusion proteins, called base editors. In base editing, a partially nuclease-inactive nickase-cas9(nCas9) protein is fused to either the cytidine deaminase APOBEC1A to enable C-G to T-A base pair changes or to an evolved TadA heterodimer to facilitate the opposite reaction, turning A-T base pairs into G-C base pairs (Gaudelli et al., 2017; Komor et al., 2016). Base editors show high efficiency and infrequent unwanted DNA changes in a variety of model systems but are strictly limited to transition DNA substitutions (Geurts et al., 2020; Pavlov et al., 2019; Zuo et al., 2019). To overcome these limitations, prime editing has been developed to enable both transition and transversion reactions as well as insertions and deletions of up to 80 nucleotides in length without the need to generate DSBs (Anzalone et al., 2019). In prime editing an nCas9 is fused to an engineered reverse transcriptase (RT) that is used to generate complementary DNA from an RNA template (PE2) (**Figure 1A**). This fusion protein is combined with a prime editing guide RNA (pegRNA) that guides the nCas9 to its target and contains the RNA template that encodes the desired edit. Upon target recognition the PAM-containing strand is nicked and the pegRNA extension binds to the nicked strand at the primer binding site (PBS). The RT-domain then uses the remainder of the pegRNA to synthesize a 3’-DNA-flap containing the edit of interest. This DNA-flap is resolved by cellular DNA repair processes that can be further enhanced by inducing a proximal second nick in the opposing DNA strand, guided by a second (PE3) guide-RNA (**Figure 1A**). Prime editing holds great promise, as it can -in theory-repair 89% of all disease-causing variants (Anzalone et al., 2019). Here, we apply this approach in patient stem cells to repair mutations in the CFTR channel that cause cystic fibrosis, a mendelian disorder with high prevalence in European ancestry.

**Figure 1:**
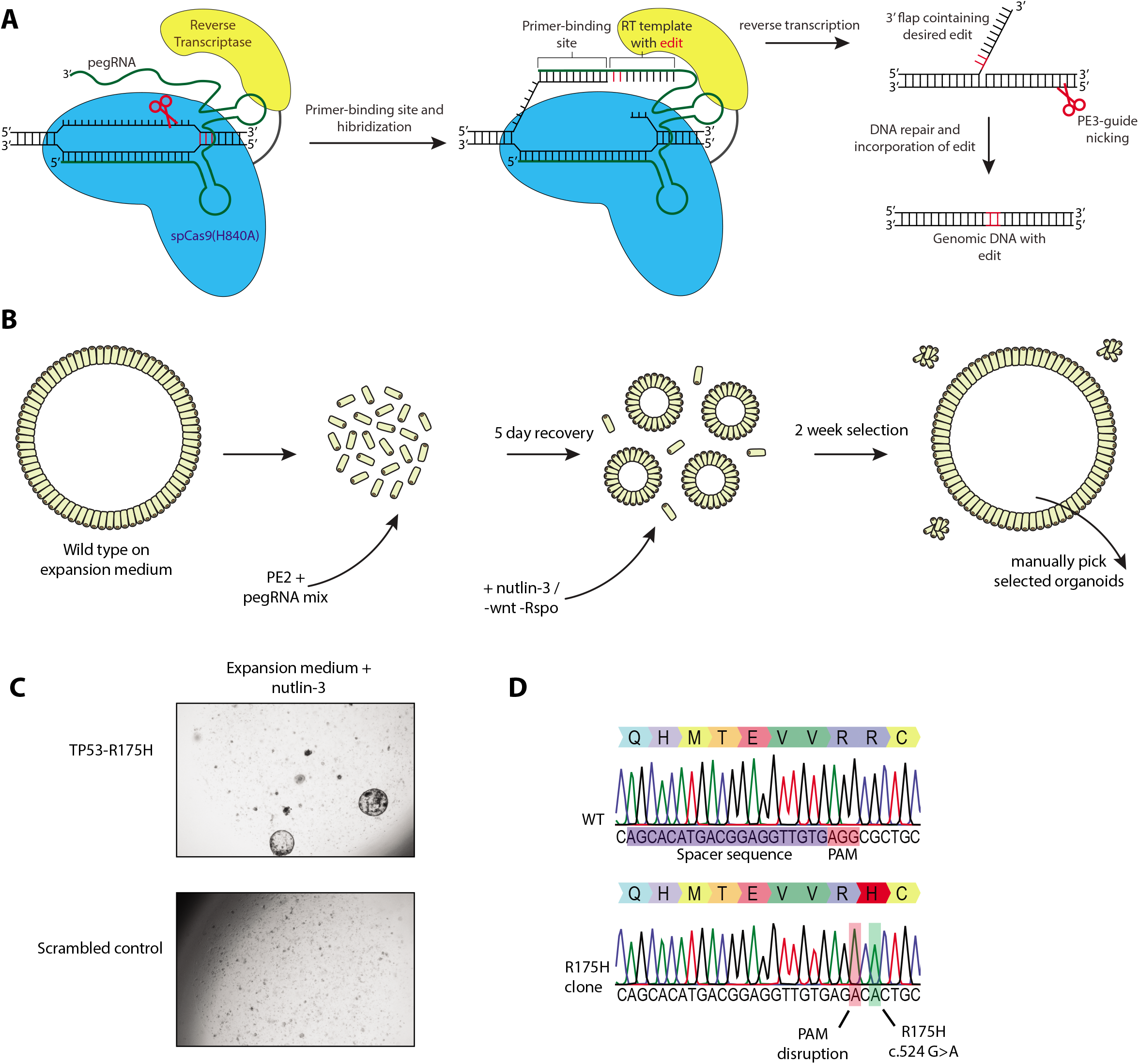
Prime editing enables generation of oncogenic mutations in intestinal organoids. (A) Principles of prime editing. Red scissors indicate nick site of the nCas9. RT = Reverse Transcriptase. (B) strategy to generate TP53 mutated human intestinal organoids. (C). Brightfield images of prime editing experiments targeting the TP53-R175H, TP53-C176F and TP53-R249S mutations compared to a negative scrambled sgRNA control. (D) Sanger sequencing trace of selected clonal organoids harbouring the TP53-R175H mutation compared to WT.

## Results

### Prime editing proof of principle in intestinal organoids

We first characterized and optimized prime editing efficacy in adult human stem cell-derived intestinal organoids by targeting TP53, a gene that is often mutated in cancer. Previously, we have shown that TP53-mutant organoids can be selected by adding nutlin-3, a molecule that inhibits the interaction between tp53 and MDM2, to the organoid culture medium (Drost et al., 2015; Matano et al., 2015). Using the pegFinder online software tool, we designed a single pegRNA and PE3-guide pair to create the R175H mutation, the most common mutation found in TP53 according to the Catalogue Of Somatic Mutations in Cancer (COSMIC) (Chow & Chen, 2020; Forbes et al., 2017). The prime editing guide RNAs were designed to integrate a protospacer adjacent motif (PAM)-disrupting mutation in order to block re-binding of Cas9 after the correct editing event has occurred. Wild-type intestinal organoids were digested into a single-cell suspension and co-transfected with the PE2 plasmid and the pegRNA/PE3-guide pair by electroporation (**Figure 1B**). Clonally selected organoids appeared after two weeks of nutlin-3 selection whereas control organoids, transfected with PE2 plasmids and a non-targeting scrambled sgRNA did not grow out (**Figure 1C**). Manual picking of selected organoids and subsequent sanger sequencing showed correct homozygous induction of the TP53-R175H(c.524 G>A) mutation in seven out of the eight clonally expanded organoids (**Figure 1D**). Next, we aimed to construct six additional mutations that are commonly found in TP53. Only the pegRNA/PE3-guide pairs designed for the induction of TP53-C176F and TP53-R249S resulted in clones capable of surviving nutlin-3 selection (**Figure 2A and D**). Manual picking of these clones followed by sanger sequencing showed correct homozygous introduction of both the C176F(c.527 G>T) and R249S mutation in the TP53(c.747 G>T) gene and included the designed PAM disruption mutation (**Figure 2B and C**). Additionally, we designed pegRNA/PE3-guide pairs to generate mutations in APC, the gene that is often the first to be mutated in colorectal cancer. Mutations in APC can be selected in culture by the removal of the expansion medium components WNT and R-spondin (**Figure 1B**) (Drost et al., 2015; Matano et al., 2015). Selection for APC mutants by removal of WNT and R-spondin after transfection of two pegRNA/PE3-guide pairs resulted in outgrowth of a single clone (**Figure 2D and E**). Interestingly, instead of the designed APC-R1450*(c.4348 C>T) mutation, sanger sequencing revealed a homozygous duplication of the 37 nucleotides directly upstream of the single stranded nick introduced by the SpCas9(H840A) (**Figure 2F**). These results indicated that prime editing can induce mutations in adult human stem cells but may yield undesired outcomes. Even though editing efficiencies varied greatly between targeted mutations, these results gave us confidence to proceed with the repair of CFTR mutations that cause cystic fibrosis.

**Figure 2:**
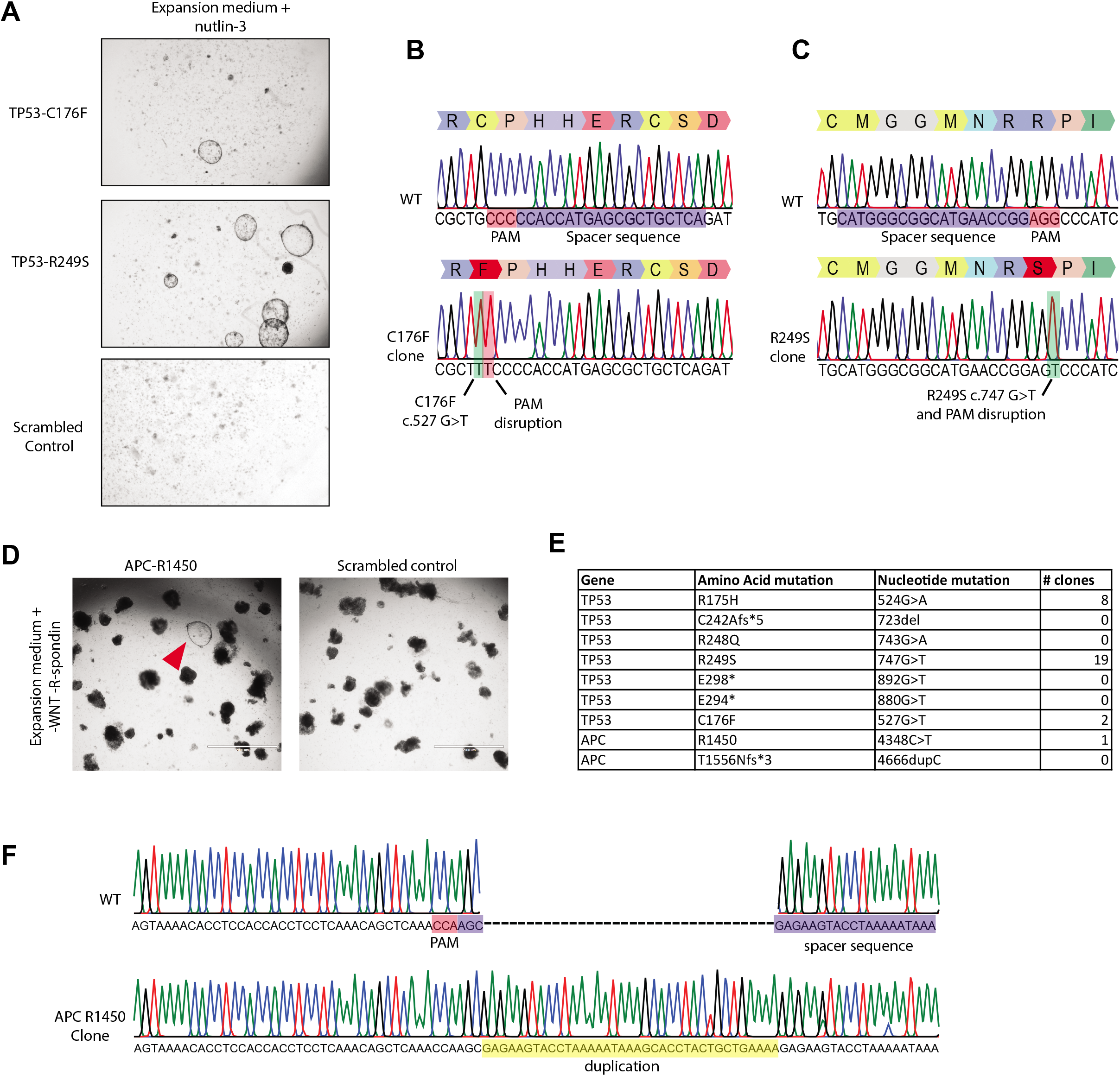
Prime editing enables generation of oncogenic mutations in intestinal organoids. (A) Brightfield images of prime editing experiments targeting the TP53-C176F and TP53-R249S mutations compared to a negative scrambled sgRNA control. (B) Sanger sequencing trace of selected clonal organoids harbouring the TP53-C176F mutation compared to WT.(C) Sanger sequencing trace of selected clonal organoids harbouring the TP53-R249S mutation compared to WT. Red box indicates PAM disruption mutation and green box indicates the desired mutation. (D) Amount of observed clones as observed after either selection with both the addition of nutlin-3 (TP53) or removal of wnt and Rspo1 (APC) from the culture medium. (E) Brightfield images of prime editing experiments targeting the APC R1450* mutation compared to a negative scrambled sgRNA control. (F) Sanger sequencing trace of selected APC R1450* clone. Insertion is shown in yellow, PAM is shown in red and spacer sequence is shown in blue.

### Repair of CFTR-F508del mutation using prime editing in intestinal organoids

Intestinal organoids are a suitable *in vitro* model of cystic fibrosis as the activity of the CFTR channel can be directly assessed by forskolin-induced intracellular cAMP levels. Wild-type organoids show a forskolin induced swelling (FIS) response whereas FIS is strongly reduced in organoids derived from people with cystic fibrosis (Dekkers et al., 2013). This assay is highly predictive for drug response and is clinically used to guide personalized treatment of CF patients in the Netherlands (Berkers et al., 2019). Previously we have shown that we can use this FIS-response as a direct functional assay for repair of the *CFTR* gene in organoids derived from CF patients, both by base editing and by classical CRISPR-mediated homology-dependent repair (HDR) (Geurts et al., 2020; Schwank et al., 2013). First, we pursued prime editing-mediated repair of the CFTR-F508del mutation (**Figure 3A**), which is the most common CFTR mutation. We transfected intestinal CF organoids carrying the homozygous CFTR-F508del mutation with a pegRNA/PE3-guide pair. Forskolin treatment two weeks after electroporation showed a swelling response in a single transfected organoid (**Figure 3B**). PCR amplification of the target site, followed by sub-cloning and sanger sequencing revealed heterozygous repair of the CFTR-F508del mutation in this selected clone (**Figure 3C**). We tried to further optimize prime editing by designing additional pegRNA/PE3-guide pairs with varying RT and PBS lengths and distance between the pegRNA and PE3-guide as these variables greatly impact editing efficiencies (Anzalone et al., 2019)(**Supplemental Figure 1A**). We compared editing efficiencies of 8 different combinations of pegRNA/PE3-guide pairs (PBS length=14 or 15 nucleotides, RT length= 17 or 37 nucleotides) directly to conventional CRISPR/Cas9-mediated HDR by counting forskolin-responsive organoids. CFTR-F508del repair by CRISPR/Cas9-mediated HDR resulted in 108 FIS responsive clones while prime editing never resulted in more than 3 repaired organoids, indicating low prime editing efficiencies at this target site (**Figure 3D**). We manually picked repaired organoids for further analysis. Repaired clonal organoid lines generated by prime editing and CRISPR/Cas9-mediated HDR exhibited FIS at WT levels or higher, indicating complete functional repair of CFTR function in these organoids. Unrepaired clones did not respond to forskolin (**Figure 3E, F and G**). Sanger sequencing followed by deconvolution of the Sanger traces of 2 additional prime edited clones and one clone repaired by CRISPR/Cas9-mediated HDR showed correct heterozygous repair of the mutation in one prime editing clone. However, the second clone as well as the clone repaired by HDR contained a small indel at the repair site in the second allele (**Supplemental Figure 1B**). These results indicated that, even though efficiencies are low and undesired outcomes may occur, prime editing can repair the CFTR-F508del mutation in patient-derived intestinal organoids.

**Figure 3:**
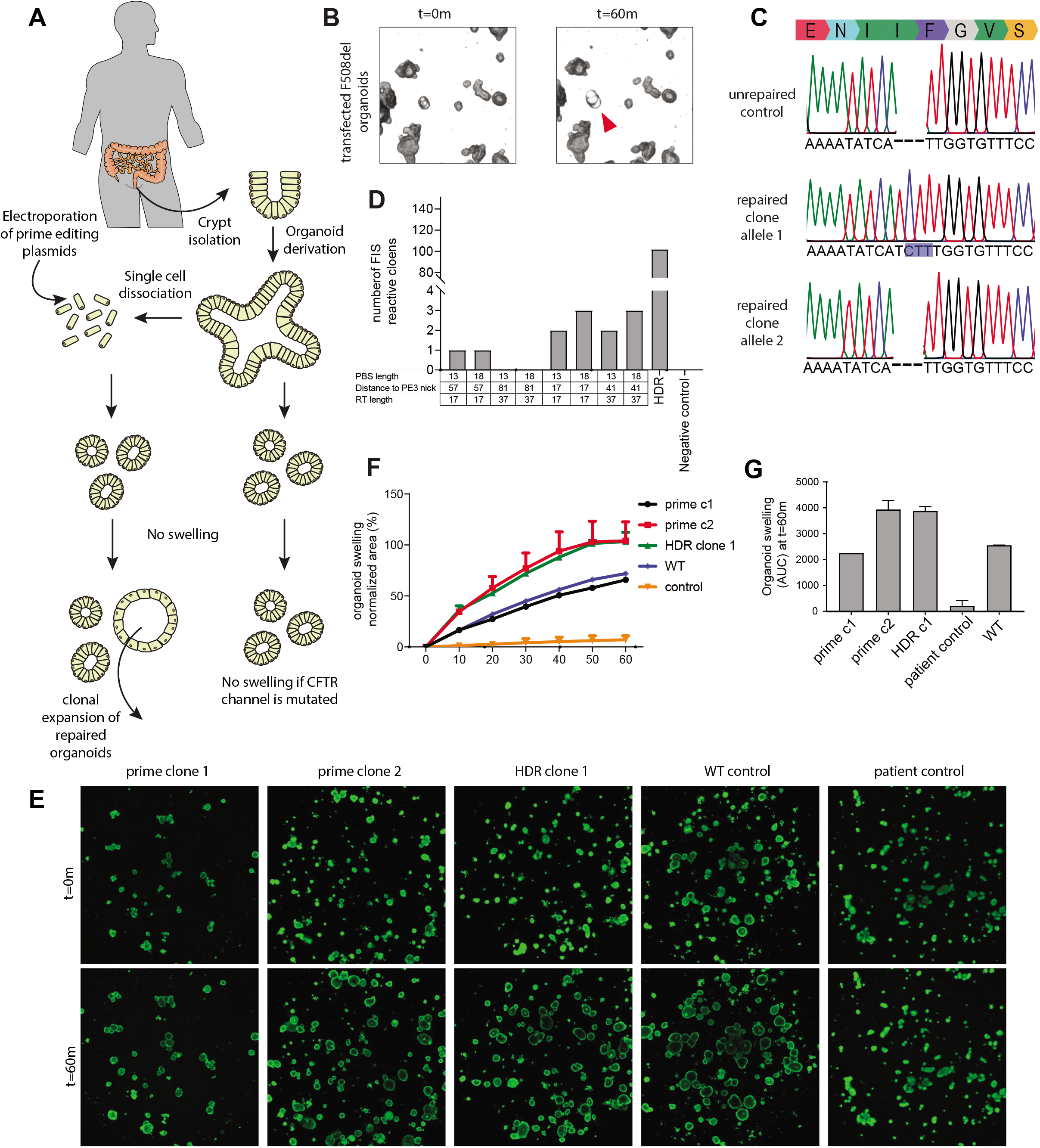
Functional repair of the CFTR-F508del mutation in patient derived intestinal organoids. (A) Experimental design of prime editing-mediated repair of CFTR mutations in human intestinal organoids. (B) Transfected CFTR-F508del organoids before (t=0) and after (t=60m) addition of forskolin. Functionally repaired organoid indicated with red arrow. (C) Sanger sequencing traces of both alleles of a functionally selected CFTR-F508del organoid line compared to unrepaired control organoids. Blue box shows the prime editing induced insertion. (D) Prime editing efficiencies for the repair of the CFTR-F508 del mutation as measured by FIS reactive organoids compared to CRISPR/Cas9-mediated HDR and a negative scrambled sgRNA control. (E) Confocal images of calcein-green-stained patient-derived intestinal organoids before and after 60 min. stimulation with forskolin (scale bars, 200 μm). (F) Per well the total organoid area (xy plane in μm2) increase relative to t = 0 (set to 100%) of forskolin treatment was quantified (n = 3). (G) FIS as the absolute area under the curve (AUC) (t = 60 min; baseline, 100%), mean ± SD; n = 3, *p < 0.001, compared to the corrected organoid clones and the WT organoid sample.

### Comparison of CFTR-R785* repair by prime editing versus repair by base editing

To directly compare prime editing to base editing, we focused on the repair of the CFTR-R785* mutation. Previously we have shown that this mutation is reparable in patient-derived intestinal organoids with an editing efficiency of ~9% while HDR efficiency was below 2% (Geurts et al., 2020). We designed two pairs of pegRNA/PE3-guides with varying PBS (13 or 18) and RT (27 or 30) lengths to find optimal prime editing conditions (**Supplemental Figure 2A**). We compared these prime editing guide pairs directly by electroporating intestinal organoids derived from a CF patient harboring the CFTR-R785* mutation on both alleles. Two weeks after transfection, forskolin-responsive clones were counted. All prime editing conditions resulted in FIS responsive clones, although editing efficiencies differed greatly (between 1 and 55 repaired clones) (**Figure 4A**). We then compared these editing efficiencies with base editing and CRISPR/Cas9-mediated HDR, using previously established reagents (Geurts et al., 2020). Forskolin treatment revealed 307 corrected organoids by ABE and 42 by CRISPR/Cas9-mediated HDR (**Figure 4A**). Repaired clonally expanded organoid lines generated by prime editing and base editing exhibited a forskolin response similar to WT levels, indicating functional repair of CFTR function in these organoids (**Figure 4B, C and D**). Sanger sequencing of three repaired organoid lines showed that two out of three clones repaired by prime editing and the ABE clone underwent correct repair of the CFTR-R785* mutation on a single allele while the second allele remained undamaged (**Figure 4E and Supplemental Figure 2B**). The third clone as well as the clone repaired by HDR contained a small indel at the repair site on the second allele indicating DNA damage (**Supplemental Figure 2B**). These results show that the current version of adenine base editing is superior to prime editing in both safety and efficiency if the mutation is targetable by adenine base editing (Geurts et al., 2020).

**Figure 4:**
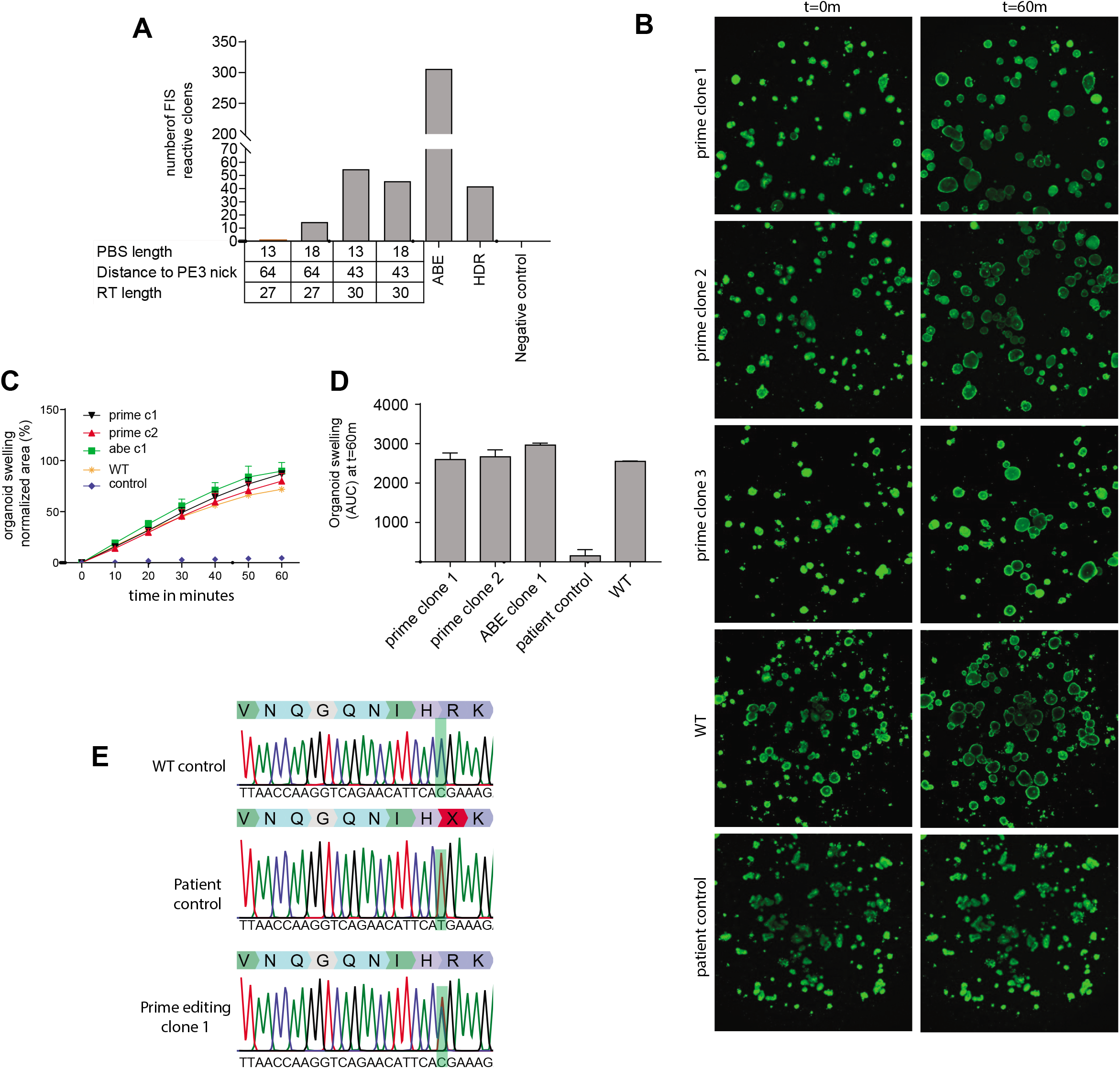
Functional repair of the CFTR-R785* mutation in patient derived intestinal organoids. (A) Prime editing efficiencies for the repair of the CFTR-R785* mutation as measured by FIS reactive organoids compared to adenine base editing, CRISPR/Cas9-mediated HDR and a negative scrambled sgRNA control. (B) Confocal images of calcein-green-stained patient-derived intestinal organoids before and after 60 min. stimulation with forskolin (scale bars, 200 μm). (C) Per well the total organoid area (xy plane in μm2) increase relative to t = 0 (set to 100%) of forskolin treatment was quantified (n = 3). (D) FIS as the absolute area under the curve (AUC) (t = 60 min; baseline, 100%), mean ± SD; n = 3, *p < 0.001, compared to the corrected organoid clones and the

## Discussion

In this study we explore the use of prime editing for functional repair of mutations in human intestinal stem cells from CF patients. Even though correct integration of the desired edits was achieved on a variety of targets, we also uncovered undesired edits, as has been detected before in mice (Aida et al., 2020). Unintended indel formation was often seen on one allele, and sometimes even on both alleles around the target site. This may be explained by the need to generate a second nick on the opposing strand close to the initial nick by the PE2 machinery. The use of two sgRNAs that nick opposing strands is known to generate indels and is even often used to increase specificity of CRISPR/Cas9-mediated genome engineering (Ran et al., 2013). Further optimization of the prime editing fusion protein may aim to render the generation of a second nick unnecessary. Nevertheless, prime editing efficiencies remained low, possibly comparable to CRISPR/Cas9-mediated HDR. Over the past years, base editing plasmids have undergone several rounds of optimization turning them into efficient genome editors (Koblan et al., 2018; Zafra et al., 2018). In our hands, base editors are superior, both in terms of efficiency and of specificity in generating only the desired mutation (Geurts et al., 2020). However, if the desired edit cannot be generated by a base editor, for instance if it regards an indel or non-transition base change, prime editing is a valuable alternative to CRISPR/Cas9-mediated HDR. Thus, prime editing is a versatile tool that can be used for disease modeling and clinical repair of most types of disease-causing mutations in human adult stem cells yet will require further improvement to allow widespread use as a technique for mutational modeling and for gene repair.

## Material and Methods

### Organoid culture

Intestinal organoids are cultured as previously described (Sato et al., 2011). In short, the wild-type human colon organoid line P26n, as previously described in (Van De Wetering et al., 2015) was cultured in domes of Cultrex Pathclear Reduced Growth Factor Basement Membrane Extract (BME) (3533-001, Amsbio). Domes were covered by medium containing Advanced DMEM/F12 (Gibco), 1x Glutamax, 10mmol/l HEPES, 100uU/ml penicillin-stroptomycin and 1x B27 (All supplied by ThermoFisher scientific), 1.25mM N-acetylcysteine, 10μM nicotinamide, 10μM p38 inhibitor SB202190 (supplied by Sigma-Aldrich). This medium was supplemented with the following growth factors: 0.4nM Wnt surrogate-Fc Fusion protein, 2% Noggin conditioned medium (U-Protein express), 20% Rspo1 conditioned medium (in-house), 50ng.ml EGF (Peprotech), 0.5 μM A83-01 and 1μM PGE2 (Tocris). Intestinal organoids derived from people with cystic fibrosis are part of a large biobank at Hub for organoids (HUB), are stored in liquid nitrogen and are passaged at least 4 times prior to electroporation experiments. Cystic fibrosis organoids are kept in Matrigel (Corning) instead of BME. Furthermore 2% Noggin conditioned medium (U-protein expess) is replaced by 10% Noggin conditioned medium (in-house). Moreover, PGE2 is excluded and 30μM of P38 inhibitor SB202190 (Sigma-Aldrich) is added to expansion medium for cystic fibrosis organoids. All organoids were passaged and split once a week 1:6 and filtered through a 40 micron cell strainer (ThermoFisher scientific) to remove differentiated structures from the culture.

### Plasmid construction

Human codon optimized prime editing constructs were a kind gift from David Liu; pCMV_PE2_P2A_GFP (Addgene plasmid #132776), pU6-pegRNA-GG-acceptor (Addgene plasmid #132777). Human codon optimized base editing constructs were a kind gift from David Liu; pCMV_ABEmax_P2A_GFP (Addgene plasmid # 112101). The empty sgRNA plasmid backbone was a kind gift from Keith Joung (BPK1520, Addgene plasmid #65777). The SpCas9 expressing vector was created by using Q5 high fidelity polymerase (NEB) to PCR amplify the Cas9-P2A-GFP cassette from pSpCas9 (BB)-2A-GFP (PX458), a kind gift from Feng Zhang (Addgene plasmid # 48138). This Cas9-P2A-GFP cassette was then cloned into the PE2 expression vector NEBbuilder HIFI assembly mastermix according to manufacturer protocols (NEB). pegRNA were created as previously described (Anzalone et al., 2019). In brief, the pU6-pegRNA-GG-acceptor plasmid was digested overnight using BsaI-HFv2 (NEB), loaded on a gel and the 2.2kb band was extracted using the QIAquick Gel extraction kit. Oligonucleotide duplexes for the spacer, scaffold and 3 ‘ -extension with their appropriate overhangs were annealed and cloned into the digested pUF-pegRNA-GG-acceptor by golden gate assembly according to the previously described protocol (Anzalone et al., 2019). PE3-guides and guides for both base editing and HDR experiments were cloned using inverse PCR together using BPK1520 as template and Q5 High fidelity polymerase. Upon PCR cleanup (Qiaquick PCR purification kit), amplicons were ligated using T4 ligase and Dpn1(both NEB) to get rid of template DNA. All transformations in this study were perfomed using OneShot Mach1t1(ThermoFisher scientific) cells and plasmid identity was checked by Sanger sequencing (Macrogen). All constructed guide-RNA sequences can be found in **Supplementary table 1**.

### Organoid electroporation

Organoid electroporation was performed with slight modifications to this previously described protocol (Fujii et al., 2015; Geurts et al., 2020). Wild type colon and intestinal organoids derived from cystic fibrosis patients were maintained in their respective expansion medium up until two days before electroporation. Two days in advance, the expansion medium was switched to electroporation medium which does not contain the growth-factors wnt and Rspo1. Rspondin-conditioned medium was replaced by Advanced DMEM-F12(Gibco) supplemented by 1x Glutamax, 10mmol/l HEPES and 100uU/ml penicillin-streptomycin (ThermoFisher scientific). Furthermore, the GSK-3 inhibitor CHIR99021(Sigma-Aldrich) was added to the medium for wnt-pathway activation and rho-kinase inhibitor Y-27632 (abmole bioscience) was added to inhibit anoikis. One day prior to electroporation 1.25%(v/v) DMSO was added to the organoid medium. On the day of electroporation the organoids were dissociated into single cells using TrypLE (Gibco) supplemented with Y-27632 at 37°C for 15 minutes. During the single-cell dissociation, the organoid suspension was vigorously pipetted every 5 minutes to keep the solution homogenous. 10^6^ cell per electroporation were resuspended in BTXpress solution and combined with 10μl plasmid solution containing 7.5 μg pCMV_PE2_P2A_GFP, pCMV_SpCas9 or pCMV_ABEmax_P2A_GFP depending on gene editing strategy and 2.5 μg per guide-RNA plasmid. In HDR experiments 2.5 μg of single stranded donor oligonucleotide, containing a WT-CFTR sequence and silent mutations to block Cas9 cleavage after repair was added to the plasmid mix. Electroporation was performed using NEPA21 with settings described before (Fujii et al., 2015). After electroporation the cells were resuspended in 600ul Matrigel or BME (50% Matrigel/BME, 50% expansion medium) and plated out in 20 μl droplet/well of a pre-warmed 48-wells tissue culture plate (Greiner). After polymerization, the droplets were immersed in 300 μL of expansion medium and the organoids were maintained at 37°C and 5% CO2.

### Phenotypic selection by forskolin induced swelling

After electroporation, organoids were expanded for 7 days and subsequently replated in 72 wells of 48-wells tissue culture plates (Greiner) to make organoids sufficiently sparse. Selection of genetically corrected organoids was based on CFTR-function restoration as assessed by adding foskolin (5μM) to the expansion medium. Pictures were made (1.25x on an EVOS FL Auto Imaging system) before and 60 minutes after forskolin addition. Organoids that showed swelling after 60 minutes were individually picked with a p200 pipette and a bend p200 pipette tip. Each individual genetically corrected organoid was dissociated into single cells using TrypLE supplemented with Y-27632 (10 μM) for 10 minutes at 37°C. The cells were plated in 20 μL matrigel droplets/picked organoid (50% matrigel, 50% CCM+) in pre-warm 48-well tissue culture plates (Greiner) and maintained at 37°C and 5% CO2.

### Phenotypic selection for oncogenic mutations

After electroporation, organoids were expanded for 5 days to offer sufficient time for recovery of the transfected cells. In prime editing experiments with the goal to mutate TP53, 10μM Nutlin-3 was added to the expansion medium. In prime editing experiments with the goal to mutate APC, both wnt-surrogate and Rspo1 were removed from the expansion medium. After two weeks individual organoids that survived selection were manually picked and clonally expanded as previously described.

### Genotyping of clonal organoid lines

Organoid DNA was harvested from 10-20μL Matrigel/BME suspension and DNA was extracted using the Zymogen Quick-DNA microprep kit. Target regions were amplified from the genome using Q5 high fidelity polymerase. Sequencing was performed using the M13F tail as all forward amplification primers for targeted sequencing contained a tail with this sequence. Prime editing, base editing and CRISPR/Cas9-mediated HDR induced genomic alterations were confirmed by Sanger sequencing (Macrogen). Subsequent sanger trace deconvolution was performed with the use of the online tool ICE by Synthego.

### FIS-Assay

To quantify CFTR function in the genetically corrected intestinal organoids, we conducted the Forskolin induced Swelling(FIS)-assay. This was done in duplicates at three independent culture time points (n=3) according to previously published protocols (Boj et al., 2017; Vonk et al., 2020). In brief, intestinal organoids were seeded in 96-well culture plates in 4μL of 50% Matrigel. Each Matrigel dome contained roughly 20-40 organoids and was immersed in expansion medium. The day after, organoids were incubated for 30 min with 3 μM calcein green (Invitrogen) to fluorescently label the organoids and stimulated with 5 μM forskolin. Every ten minutes the total calcein green labeled area per well was monitored by a Zeiss LSM800 confocal microscope, for 60 minutes while the environment was maintained at 37°C and 5% CO2. A Zen Image analysis software module (Zeiss) was used to quantify the organoid response (area under the curve measurements of relative size increase of organoids after 60 minutes forskolin stimulation, t = 0 min baseline of 100%).

## Author contributions

Conceptualization, M.H.G. and H.C.; Prime editing optimization in organoids, M.H.G, L.C and C.P.M.; Fluid secretion & Biochemical Assays E.D.P.; Cloning of Prime editing plasmids, M.H.G, L.C and A.A.R.; Functional selection of oncogenic mutations in organoids, M.H.G and M.B.; Writing original draft M.H.G and H.C.; Supervision, J.M.B and H.C.

---

**Supplementary Figure 1:**
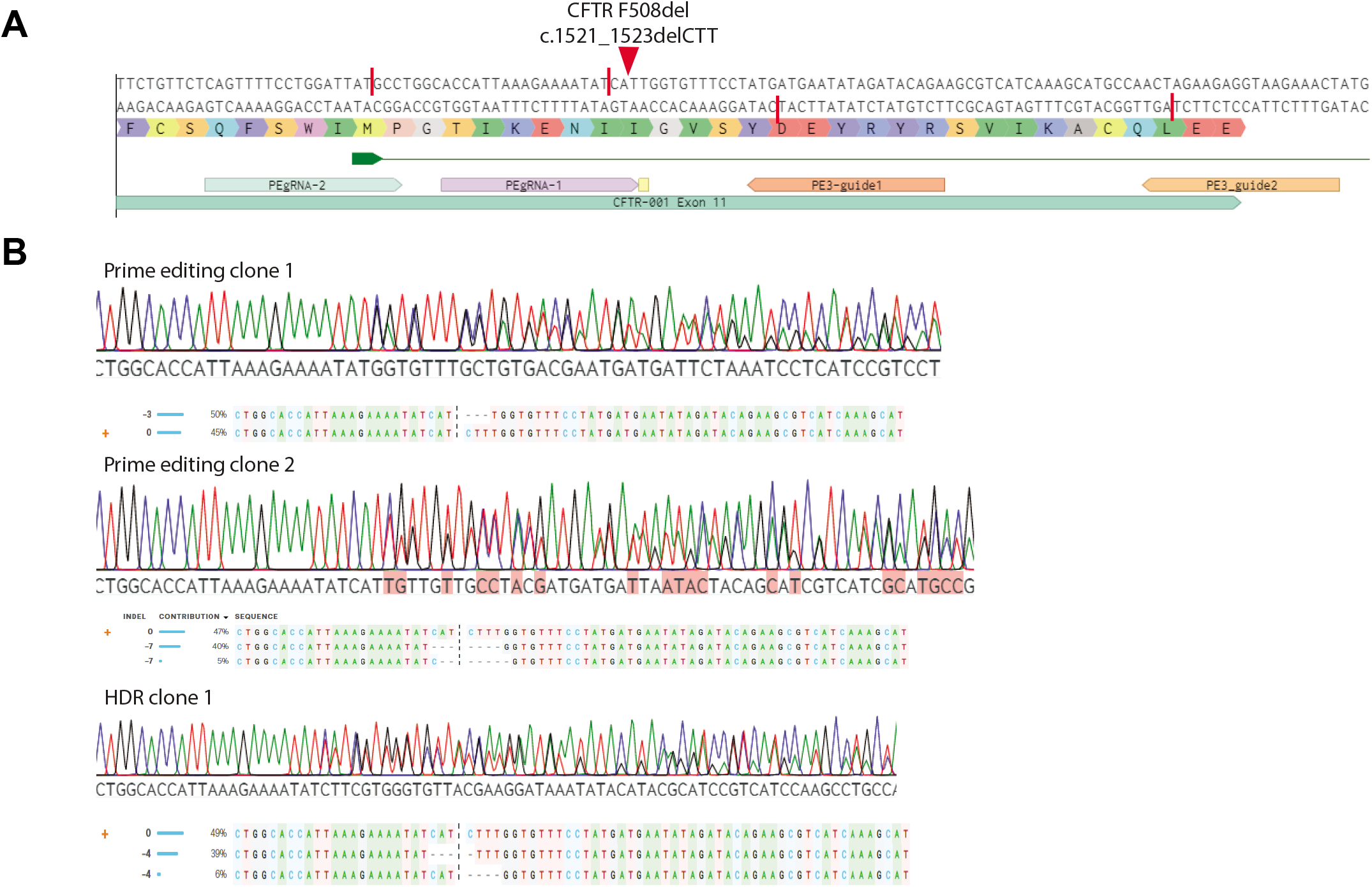
CFTR-F508del prime editing in intestinal organoids. (A) guide-RNA design for the repair of the CFTR-F508del mutation in human intestinal organoids. Red bars show the nickase sites of the guide sequences and the red arrow shows the mutation site in the DNA of organoids derived from a person with cystic fibrosis. (B) Sanger sequencing traces and deconvoluted alleles of 2 additional prime editing clones and one HDR clone that had been selected for by FIS after transfection.

**Supplementary Figure 2:**
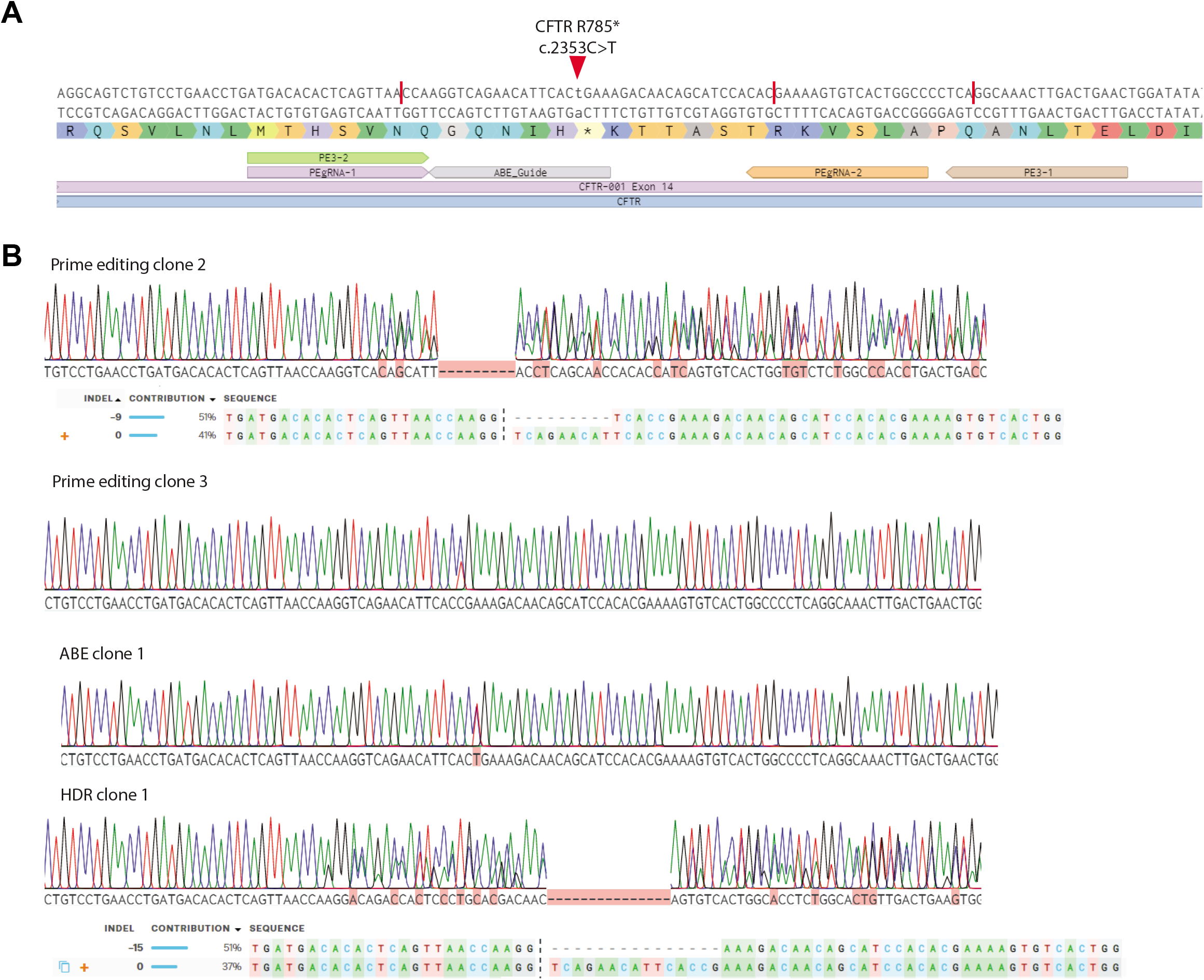
CFTR-R785* prime editing in intestinal organoids. (A) guide-RNA design for the repair of the CFTR-R785* mutation in human intestinal organoids. Red bars show the nickase sites of the guide sequences and the red arrow shows the mutation site in the DNA of organoids derived from a person with cystic fibrosis. (B) Sanger sequencing traces and deconvoluted alleles of 2 additional prime editing clones,one HDR clone and one clone repaired by adenine base editing that had been selected for by FIS after transfection.

**Supplementary Table 1:**
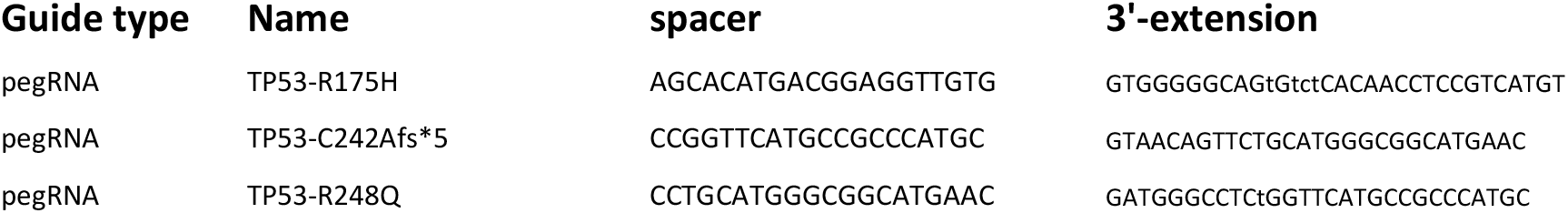
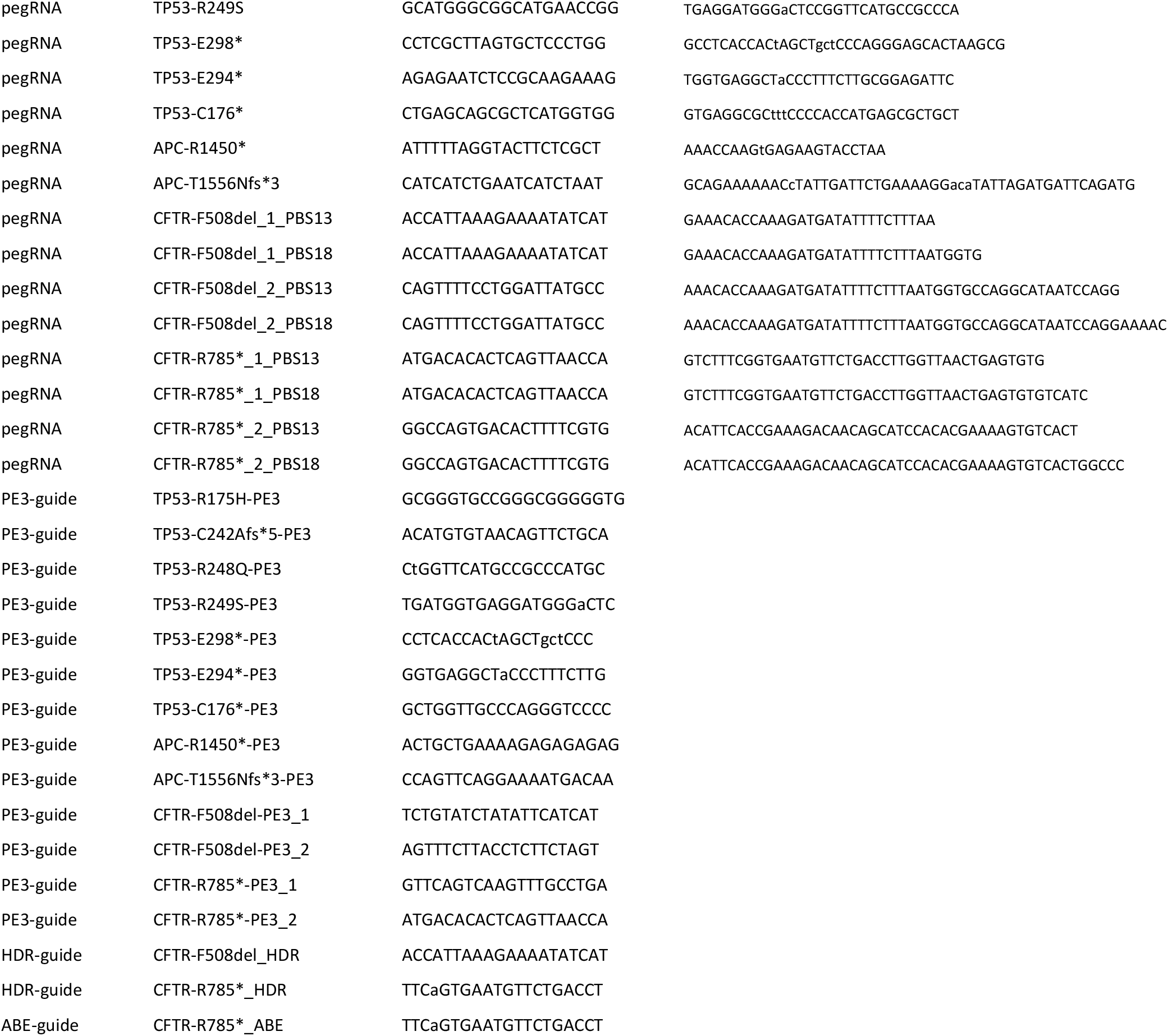
guide-RNA’s used in this study

